# The ability of human TIM1 to bind phosphatidylethanolamine enhances viral uptake and efferocytosis compared to rhesus and mouse orthologs

**DOI:** 10.1101/2024.07.29.605603

**Authors:** Lizhou Zhang, Claire E. Kitzmiller, Audrey S. Richard, Sonam Popli, Hyeryun Choe

## Abstract

T-cell Immunoglobulin and Mucin (TIM)-family proteins facilitate the clearance of apoptotic cells, are involved in immune regulation, and promote infection of enveloped viruses. These processes are frequently studied in experimental animals such as mice or rhesus macaques, but functional differences among the TIM orthologs from these species have not been described. Previously, we reported that while all three human TIM proteins bind phosphatidylserine (PS), only human TIM1 (hTIM1) binds phosphatidylethanolamine (PE), and that this PE-binding ability contributes to both phagocytic clearance of apoptotic cells and virus infection. Here we show that rhesus macaque TIM1 (rhTIM1) and mouse TIM1 (mTIM1) bind PS but not PE and that their inability to bind PE makes them less efficient than hTIM1. We also show that alteration of only two residues of mTIM1 or rhTIM1 enables them to bind both PE and PS, and that these PE-binding variants are more efficient at phagocytosis and mediating viral entry. Further, we demonstrate that the mucin domain also contributes to the binding of the virions and apoptotic cells, although it does not directly bind phospholipid. Interestingly, contribution of the hTIM1 mucin domain is more pronounced in the presence of a PE-binding head domain. These results demonstrate that rhTIM1 and mTIM1 are inherently less functional than hTIM1, owing to their inability to bind PE and their less functional mucin domains. They also imply that mouse and macaque models underestimate the activity of hTIM1.

**SIGNIFICANCE:** We previously reported that human T-cell Immunoglobulin and Mucin protein 1 (TIM1) binds phosphatidylethanolamine (PE) as well as phosphatidylserine (PS) and that PE is exposed on the apoptotic cells and viral envelopes. Moreover, TIM1 recognition of PE contributes to phagocytic clearance of apoptotic cells and virus uptake. Here we report that unlike human TIM1, murine and rhesus TIM1 orthologs bind only PS, and as a result, their ability to clear apoptotic cells or promote virus infection is less efficient. These findings are significant because they imply that the activity of TIM1 in humans is greater than what the studies conducted in common animal models would indicate.

## INTRODUCTION

T-cell Immunoglobulin Mucin domain (TIM)-family proteins are a group of cell-surface receptors that recognize phosphatidylserine (PS) exposed on apoptotic cells and initiate phagocytic clearance of those cells, namely, efferocytosis (1, 2). TIM proteins are glycoproteins consisting of four major domains: an immunoglobulin variable-like N-terminal globular domain (IgV), a heavily O-glycosylated stalk-like mucin domain, a transmembrane domain, and a cytoplasmic domain (Fig. 1A). Of these, the IgV head domain contains a binding site for PS (3, 4). Generally restricted to the cytosolic leaflet of the plasma membrane bilayer, PS flips to the outer leaflet upon the onset of apoptosis, where it acts as an “eat-me” signal for phagocytes (5, 6). Human TIM family consists of three members (hTIM1, hTIM3, and hTIM4). All three members bind PS and mediate efferocytosis (4, 7–10). TIM4 carries out this role on the professional phagocytes such as dendritic cells and macrophages (7, 11). TIM1 is expressed on a subset of T and B cells (12–15), but expression can also be induced on various epithelial cells, including those in the lung, kidney, mammary gland, retina, placenta, and testis, and assumes its role of phagocytic clearing of neighboring cells when they undergo apoptosis (10, 13, 16–24).

**Fig. 1.**
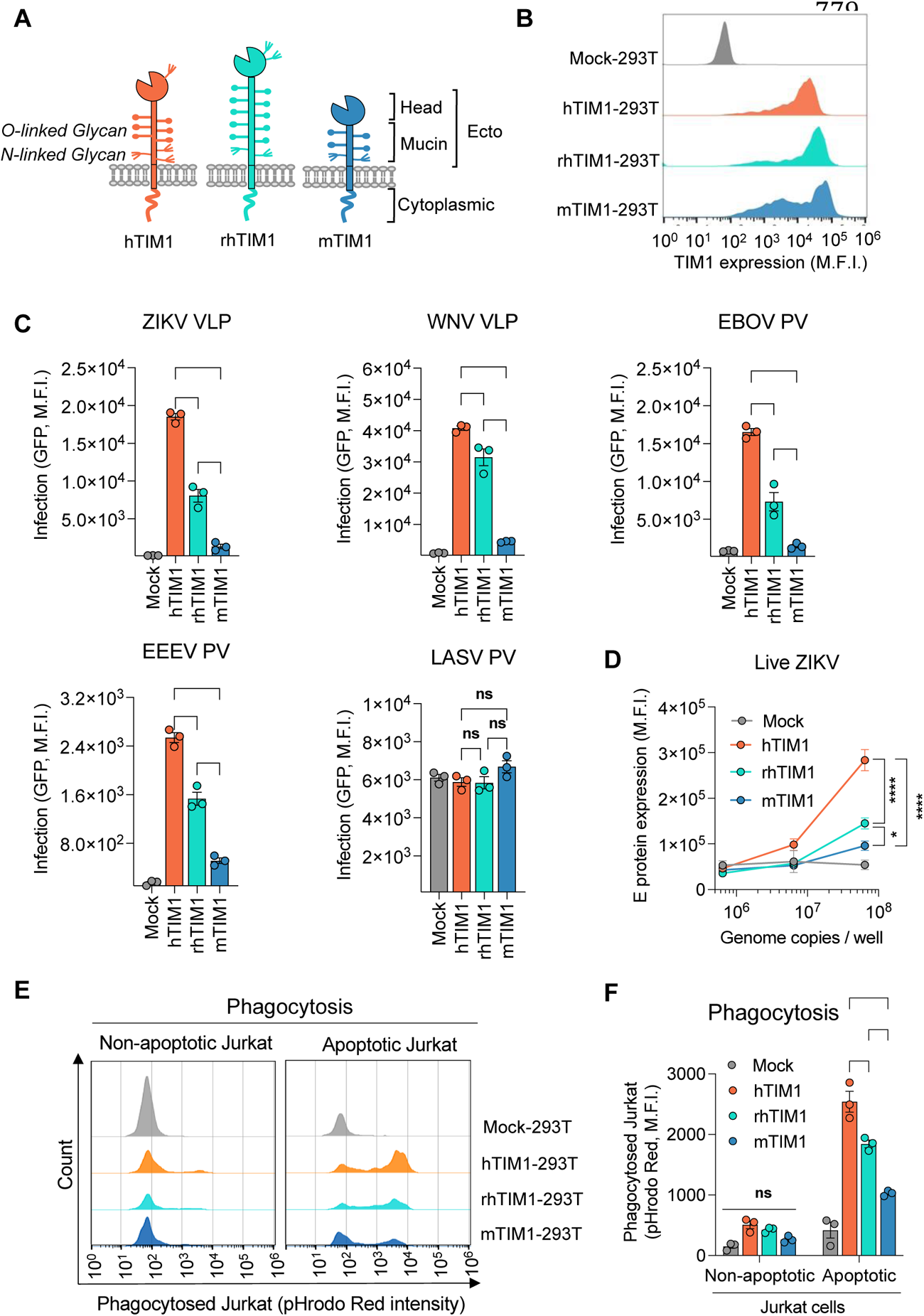
mTIM1 and rhTIM1 are not as efficient as human TIM1 in mediating virus infection or efferocytosis. **(A)** Schematic diagram of human, rhesus, and mouse TIM1 orthologs. **(B)** Cell surface expression of TIM1 orthologs in stable 293T cells. Cells were stained with anti-MYC antibody as all TIM1 molecules were MYC-tagged at their N-terminus. Mock-293T cells were produced in the same way as TIM1 stable cells except for TIM1 expression. **(C)** Infection of Mock-or TIM1-293T stable cells with VLPs or PVs. Cells were infected with VLPs (ZIKV or WNV) or PVs (EBOV, EEEV, or LASV) at 37°C for 1 h and analyzed for GFP expression 24 hours later. Each dot in the graph represents one independent experiment. M.F.I., mean fluorescence intensity. **(D)** Infection of Mock- or TIM1-293T stable cells by live ZIKV. Cells were infected with the indicated amounts of replication-competent ZIKV. One day later, ZIKV E protein was detected with the 4G2 antibody in permeabilized cells. **(E)** Phagocytosis of apoptotic Jurkat cells by Mock- or TIM1-293T cells. Jurkat cells were treated with 1 μM Actinomycin D or DMSO (control) for 15 hours in a CO_2_ incubator to induce apoptosis (see Fig. S1A), loaded with 0.1 μM pHrodo Red dye at 37°C for 1 h, and washed before incubation with Mock- or TIM1-293T cells at a 10:1 ratio of Jurkat to 293T cells. After 1 hour of co-incubation, unbound Jurkat cells were removed by PBS wash and 293T cells were detached from the plate by trypsinization for flow cytometric analysis. Gating strategy is shown in Fig. S1B. Images shown are the representative of three independent experiments. To demonstrate bright red florescence is emitted from phagocyted, rather than bound but not internalized, Jurkat cells, co-incubation was also performed on ice (see Fig. S1B). **(F)** Quantification of the efferocytosis results shown in (E) and two additional assays based on M.F.I. of pHrodo Red within the 293T cell gate. (C, D, F) Data are presented as Mean ± SEM. Statistical significance was analyzed by One-way ANOVA for (C) or by Two-way ANOVA for (D) and (F). **p < 0.01, ***p < 0.001, and ****p < 0.0001; ns, not significant.

Whereas all three members of the human TIM family bind PS (4, 7–10), only hTIM1 additionally binds phosphatidylethanolamine (PE), which we and others showed previously (10, 25). Like PS, PE is also restricted to the inner leaflet of the plasma membrane but flips to the outer leaflet during apoptosis (25–28), and we have shown that PE exposed on the surface of the apoptotic cells and virions contributes to hTIM1-mediated efferocytosis and virus entry, respectively (25).

Because efficient clearance of apoptotic cells is essential for the maintenance of healthy tissues and immunity, failure to detect apoptotic signals is associated with altered immune tolerance and autoimmunity. Thus, like other PS-binding molecules involved in the clearance of apoptotic cells, TIM1 is implicated in immune regulation, inflammation control, and autoimmune diseases (1, 18, 29–34). TIM1 was also identified as an asthma-susceptibility gene (35) and kidney injury molecule, KIM-1 (10), and is alternatively known as HAVCR1 after it was reported as a receptor for hepatitis A virus (36). In addition, it is well established that a wide range of enveloped viruses efficiently utilize TIM proteins to infect cells (37–42). This mechanism is known as “apoptotic mimicry” (43–45), in which viruses enter cells by disguising as apoptotic bodies and taking advantage of the signals for endocytosis and immune suppression transmitted by TIM family members (37–42) or other PS-binding molecules (37, 46–49).

Because of the important roles of TIM1 in immune regulation, autoimmune diseases, and virus infection, multiple TIM1 knock-out mice were generated to study its roles *in vivo* (15, 23, 50, 51). Mouse TIM (mTIM) family consists of eight members. Based on sequence, functional, and structural data, mTIM1, 3, and 4 are considered orthologs of hTIM1, 3, and 4, respectively (1). The ligand specificity of these mouse TIM molecules, however, is less well characterized than that of human counterparts. Further, no information is available for the TIM orthologs of rhesus macaque (rhTIM), another important experimental animal species.

Mutant mice, in which TIM1 mucin domain is deleted, developed autoimmune diseases, exhibited defects in regulatory B cell function (15), and were shown to be defective in efferocytosis in the kidney tubules (21), demonstrating the importance of the mucin domain in TIM1 function. In addition, the mucin domain of hTIM1 was shown to be associated with asthma (34, 52). The mucin domains of TIM1 from different species vary in their length and glycosylation (Fig. 1A). Whether the differences in the mucin domain among these TIM1 orthologs contribute to efferocytosis or virus infection is not known.

We show here that mTIM1 and rhTIM1 are less efficient than hTIM1 in mediating efferocytosis and entry of retroviral pseudoviruses (PVs) of ebolavirus (EBOV) and eastern equine encephalitis virus (EEEV) as well as virus-like particles (VLPs) of Zika virus (ZIKV) and West Nile virus (WNV). They are also less active than hTIM1 in mediating the infection of live ZIKV. We demonstrate that the reason for their lower efficiency is because rhTIM1 and mTIM1 bind only PS whereas hTIM1 can bind PS and PE. We further demonstrate that alteration of only two residues in the head domain of mTIM1 or rhTIM1 enables them to bind PE as well as PS and enhances their ability to mediate virus entry and efferocytosis. In addition, we show that a mTIM1 variant whose mucin domain is replaced with that of hTIM1 exhibits higher efficiency in mediating efferocytosis and virus infection but only when the human mucin domain is combined with the PE-binding mutation in the head domain. These results inform the differences in ligand specificity among TIM1 orthologs and imply that TIM1 functions assessed in mouse or rhesus macaque models likely underrepresent those in human.

## RESULTS

### mTIM1 and rhTIM1 are not as efficient as hTIM1 in mediating virus infection or efferocytosis

Because mouse and rhesus macaque are the two most frequently used animal species for *in vivo* studies, we compared the efficiency of mTIM1 and rhTIM1 to that of hTIM1 for their well-established functions: mediating efferocytosis and virus entry. We first investigated their ability in supporting virus entry using ZIKV and WNV VLPs and EBOV and EEEV PVs. We previously observed that flaviviruses and flavivirus VLPs were one of the most avid users of hTIM1 as an entry factor (25, 37, 39). We also previously observed that Lassa fever virus (LASV) PV did not efficiently utilize hTIM1 to enter HEK293T or NIH3T3 cells (39), which is likely because of the presence of its high-affinity receptor, alpha-dystroglycan (53), in those cells (25). Thus, LASV PV was included as a negative control. HEK293T cells stably expressing hTIM1, rhTIM1, or mTIM1 (hTIM1-293T, rhTIM1-293T, and mTIM1-293T, respectively) with the MYC tag at their N-terminus, were generated. Although the expression levels of the three TIM1 orthologs were comparable, mTIM1 expression was the highest with that of hTIM1 the lowest (Fig. 1B). These cells were infected with the indicated VLPs or PVs encoding enhanced green fluorescent protein (eGFP). As Fig. 1C shows WNV and ZIKV VLPs, and EBOV and EEEV PVs, entered hTIM1-293T cells with much higher efficiency than they entered rhTIM1- or mTIM1-293T cells. As expected, LASV PV entry was not affected by the expression of any TIM1 molecule. We then confirmed these results, using live ZIKV. We infected the three stable TIM1-293T cells with varying amounts of replication-competent ZIKV, and the infection level was assessed by staining the E protein in the permeabilized cells with the pan-flavivirus antibody, 4G2 (Fig. 1D). Like its VLP, live ZIKV more efficiently infected hTIM1-293T cells than rhTIM1- or mTIM1-293T cells. We next compared the ability of these TIM1 orthologs to mediate efferocytosis. Jurkat cells, a human cell line derived from T-cell leukemia, were treated with Actinomycin D to induce apoptosis, and approximately 85% of these cells were apoptotic, indicated by Annexin V staining (Fig. S1A). Apoptotic Jurkat cells were loaded with pHrodo Red and incubated with hTIM1-, rhTIM1-, or mTIM1-HEK293T cells to measure phagocytic uptake. Control Jurkat cells were treated with DMSO. pHrodo Red is faintly fluorescent at neutral pH, but its fluorescence is substantially enhanced in an acidic environment such as inside the phagosomes (54). After 1 hour of incubation at 37°C, unbound Jurkat cells were removed. To prevent activation of pHrodo Red, phosphate buffered saline (PBS, pH 7.4), but no acidic buffer, was used to remove the attached but not internalized Jurkat cells. Although PBS washing did not completely remove unbound cells, residual or uninternalized Jurkat cells did not emit significant level of fluorescence (Fig. S1B, on ice) because as aforementioned, pHrodo Red fluorescence is weak at neutral pH. Therefore, total fluorescence from within the TIM1-293T cell gate (Fig. S1B, right panels) was analyzed as a measure for phagocytosis. As Fig. 1E and 1F show, while robust phagocytosis of apoptotic Jurkat cells was observed with all three TIM1-293T cells, significantly higher fluorescence was emitted from hTIM1-293T cells compared to mTIM1- or rhTIM1-293T cells. To prove that this fluorescence was from the phagocytosed rather than the attached but uninternalized Jurkat cells, we incubated hTIM1-293T cells with the Actinomycin D treated and pHrodo Red-loaded Jurkat cells on ice or at 37°C, and fluorescence was measured. TIM1-293T cells kept on ice, to which Jurkat cells were attached but not internalized, emitted only weak fluorescence, while the same cells incubated at 37°C emitted robust fluorescence (Fig. S1B, right panels). This result shows that the high fluorescence shown in Fig. 1E is from the phagocytosed Jurkat cells. Together, these data demonstrate that hTIM1 is more efficient than mTIM1 or rhTIM1 in mediating virus infection and efferocytosis.

### rhTIM1 and mTIM1 bind only PS, while hTIM1 binds PE as well as PS

To identify the features of hTIM1 that make it more efficient than mTIM1 and rhTIM1, we examined whether these three TIM1 orthologs were able to bind the phospholipid (PL) ligands with comparable efficiency. We and others previously showed that the cells undergoing apoptosis expose PE as well as PS on their surface (25–27) and that hTIM1 bound PE as efficiently as PS (10, 25).

Because PE-binding ability of mTIM1 and rhTIM1 has not been shown, we performed PL ELISA assays using the head domain of the three TIM1 orthologs. The indicated PL dissolved in methanol, was air-dried on 96-well plates. Phosphatidylcholine (PC), sphingomyelin (SPH), and phosphatidylinositol (PI) were used as negative controls. Because TIM molecules bind their PL ligands through their globular head domain (4, 7), we used the constructs in which the TIM1 head domain was fused to the Fc region of the human IgG1 (TIM1(head)-Fc) to detect PE or PS binding. To avoid the avidity effect contributed by dimerization of the Fc region, mutations L368R, F405H, and Y407E in addition to the three Cysteine mutations (C310A, C316N, and C319G) were introduced to the Fc region, generating monomeric Fc-fusion forms (Fc_mono_) (55). As Fig. 2 shows, while hTIM1(head)-Fc_mono_ binds both PE and PS equally well, mTIM1(head)-Fc_mono_ and rhTIM1(head)-Fc_mono_ bind only PS. None of TIM1 molecules binds PC, PI, or SPH. These data demonstrate a broader ligand specificity of hTIM1 compared to mTIM1 or rhTIM1 and suggest that the higher efficiency of hTIM1 in mediating virus entry and efferocytosis is, at least partially, derived from its ability to bind PE in addition to PS.

**Fig. 2.**
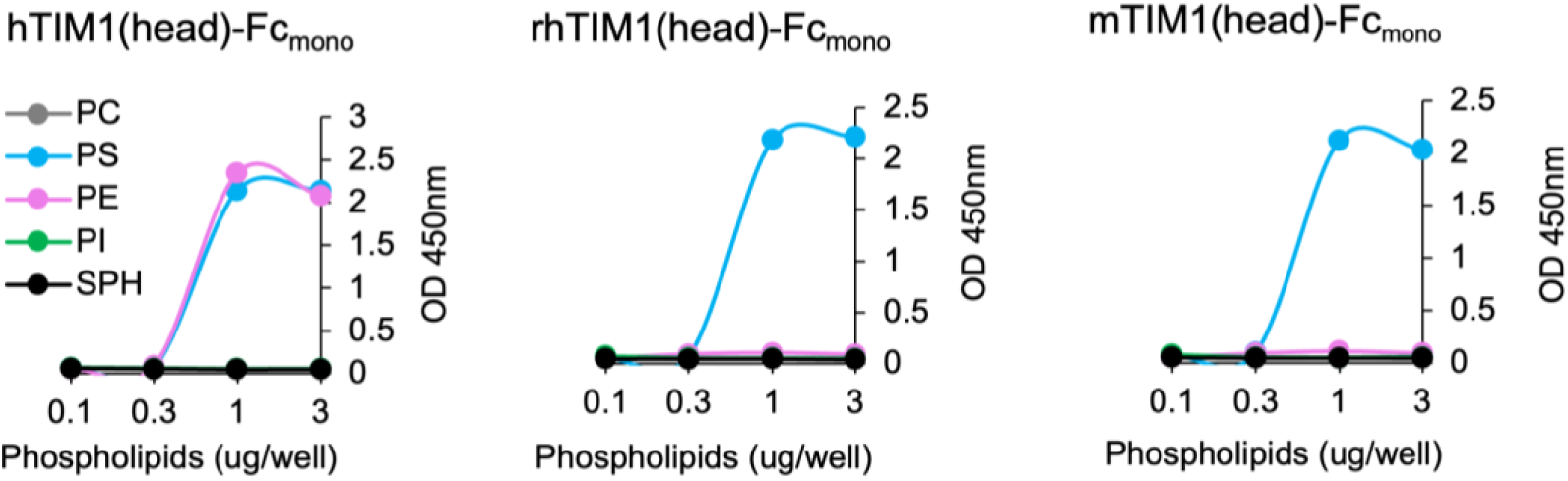
hTIM1 binds both PE and PS whereas rhTIM1 and mTIM1 bind only PS. Increasing amount (0.01 to 3 μg per well) of the indicated phospholipids dissolved in methanol was completely air dried on ELISA plates. These plates were washed with 0.05% TBST and blocked with 1% BSA and incubated for 1 h at room temperature with 100 ul of 1 nM TIM1(head)-hFc_(mono)_ in TBS containing 2 mM Ca^2+^. The data shown are the representatives of three independent experiments with similar results.

### Alteration of two residues allows rhTIM1 and mTIM1 to bind PE as well as PS

To investigate whether the PE-binding ability is the source for higher efficiency of hTIM1, we sought to modify mTIM1 and rhTIM1 to bind PE as well as PS. Structure studies show that TIM molecules are structurally conserved within the TIM family of different animal species, and that PS binds the residues located inside the cavity formed by the CC’ and FG loops (Fig. 3A and B) (3, 4). Therefore, we constructed two chimeras in which N- and C-terminal halves of hTIM1, which contains CC’ or FG loop, respectively, were swapped with those of rhTIM1 (Fig. 3B). We assessed these chimeras for their ability to bind PL ligands and observed both halves of hTIM1 are necessary for maximum PE binding (Fig. 3C).

**Fig. 3.**
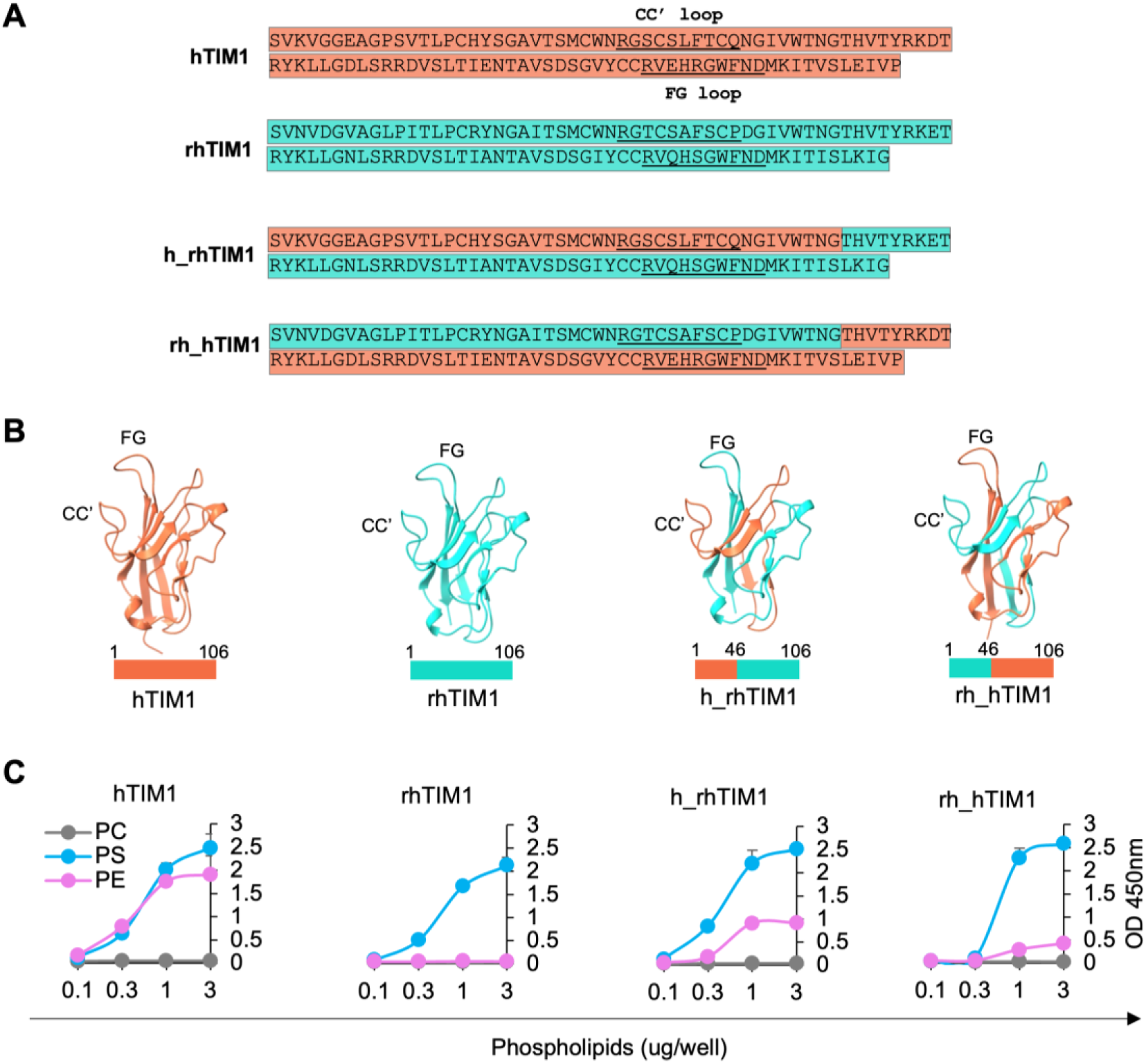
Both N- or C-terminal halves of hTIM1 are required for binding PE. **(A)** Amino acid sequences of the head domain of hTIM1, rhTIM1, and the chimeras used in this study. hTIM1 sequence is highlighted in orange, and rhTIM1 in cyan. The CC’ and FG loop sequences are underlined. **(B)** Structures of the head domain of hTIM1, rhTIM1, and their chimeras. hTIM1 structure is derived from PBD (ID: 5DZO), but others are modeled by <u>Swiss-Model</u> using 5DZO as a template. hTIM1 portion is in orange, and rhTIM1 portion in cyan. The numbers above the bars underneath the structures indicate TIM1 residues. **(C)** Phospholipid ELISA assays. Assays were performed as described in Fig. 2 legend except for the inclusion of chimeric molecules. The data shown are the representatives of three independent experiments.

Guided by the sequence differences among TIM1 molecules (Fig. 4A), structural differences between PE ad PS (Fig. 4B), and different binding modes to hTIM1 by PE and PS (Fig. 4C and D), we rationally selected several residues in the CC’ and FG loops of mTIM1 and rhTIM1 and mutated them singly or in combination. The resulting mutants were assessed for their ability to bind PE and PS. As Fig. 4E shows, in the case of rhTIM1, change of one residue each in the CC’ (A34L) and FG (Q88E) loops was necessary to gain PE-binding ability (34L88E-rhTIM1), whereas alteration of two-residues, S36L and S37F, in the CC’ loop enabled mTIM1 to bind PE as well as PS (36L37F-mTIM1).

**Fig. 4.**
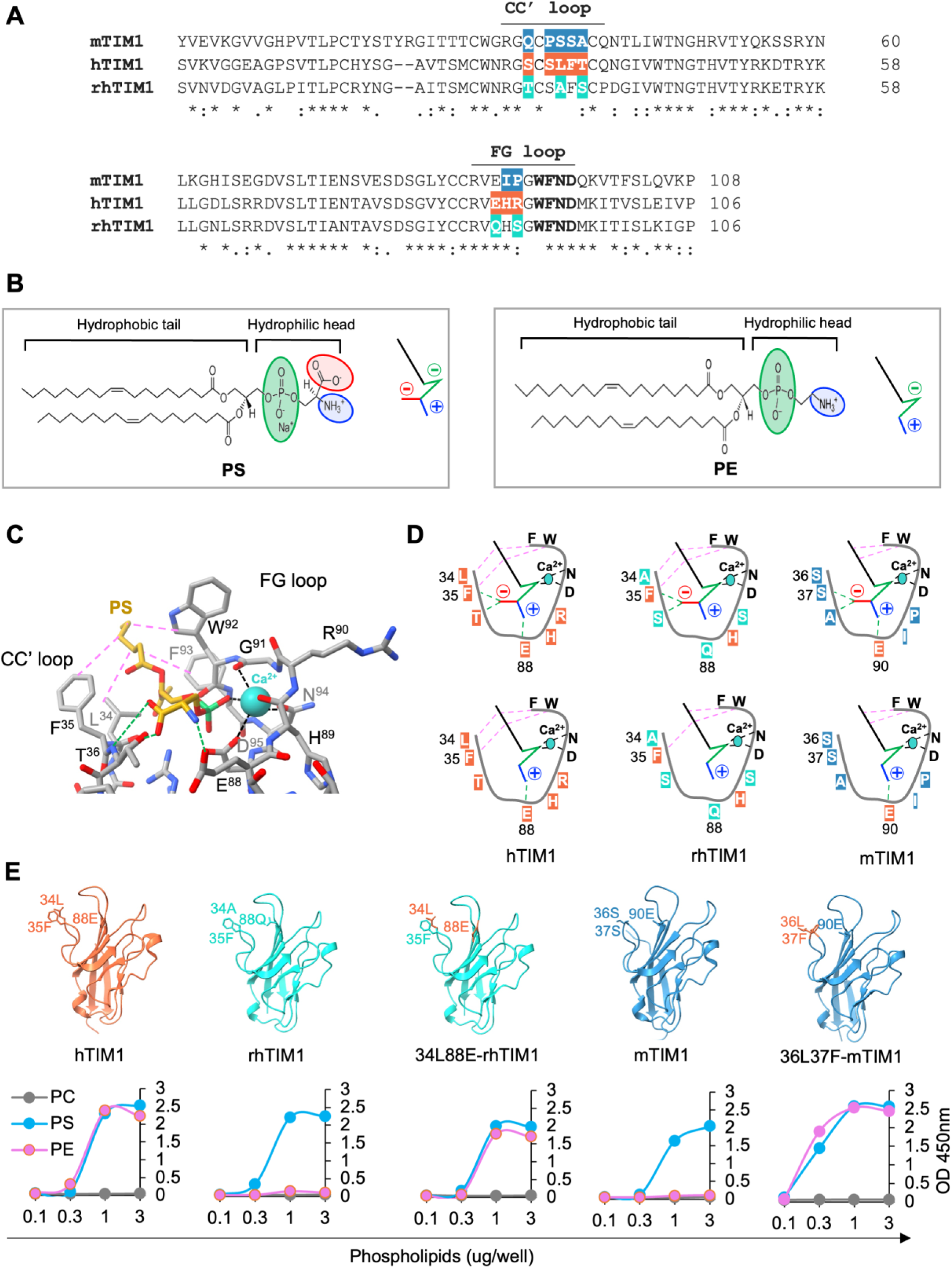
Alteration of two residues allows rhTIM1 and mTIM1 to bind PE as well as PS. **(A)** Alignment of amino acid sequences of the hTIM1, rhTIM1, and mTIM1 head domain. The CC’ and FG loops are indicated. Within the CC’ and FG loop regions, the non-conserved residues are highlighted in orange for hTIM1, in cyan for rhTIM1, and in blue for mTIM1. The bolded residues in the FG loop were reported to facilitate PS binding in the presence of Ca^2+^ (3). **(B)** Chemical structures of PE and PS are shown. Their schematic diagrams shown on the right side within the panel are used in (D). In the structures and schematic diagrams, a phosphate group is marked in green, an amine group in blue, and a carboxyl group in red. The black stick in the schematic diagrams represents the hydrophobic fatty acid tails. **(C)** Expanded view of the interface between PS and hTIM1. This structure is modeled using the PS-TIM4 complex as a template (PBD: 3BIB) and shows three major types of interactions formed between PS and hTIM1 residues: Calcium-mediated interactions are shown by black dashed lines, hydrogen bonds by green dashed lines, and hydrophobic interactions by purple dashed lines. **(D)** Schematic diagrams for the three types of interaction between the indicated TIM1 ortholog and PS (upper panels) or PE (lower panels). The same dashed lines as in (C) are used to indicate the types of interaction. **(E)** PE-binding mTIM1 and rhTIM1 mutants. The upper panel shows the location of the residues involved in binding PE. The structure of WT hTIM1 is derived from PDB: 5DZO and WT mTIM1 from PDB: 2OR8. WT and mutant rhTIM1 are modeled based on PDB: 5DZO, and mutant mTIM1 is modeled based on PDB: 2OR8. The lower panel shows PE or PS binding in phospholipid ELISA assays by WT and mutant TIM1 molecules. The data shown are the representatives of three independent experiments.

### PE-binding mutants of mTIM1 and rhTIM1 more efficiently mediate virus entry and efferocytosis

To determine whether the gained PE-binding feature of TIM1 variants correlates with functional enhancement, we examined the ability of 36L37F-mTIM1 to support ZIKV and WNV VLP entry and phagocytosis of apoptotic cells. We first generated stable cells expressing wild-type (WT) or 36L37F-mTIM1 and noticed that their expression levels were widely different in the stable cells generated through drug selection; 36L37F-mTIM1 expressed at much lower level compared to WT-mTIM1. Therefore, we conducted our studies at multiple TIM1 expression levels by transducing HEK293T cells with varying amounts of retroviral vectors expressing TIM1 molecules without drug selection. hTIM1 was used as a comparand. Next day, cells were split for VLP infection and to assess TIM1 expression. The following day, cells on 48-well plates were infected with indicated VLP, and those on 6-well plates were stained with anti-MYC antibody to measure TIM1 expression level. To assess VLP entry level, the cells infected with VLPs were analyzed for GFP expression 24 h later. When VLP entry was plotted against TIM1 expression levels, 36L37F-mTIM1 supported ZIKV and WNV VLP entry much more efficiently than did WT-mTIM1 at a wide range of expression levels (Fig. 5A and B). Two additional experiments showed nearly indistinguishable results (Fig. S2A and B). We also similarly characterized the PE-binding mutant of rhTIM1 and obtained comparable results (Fig. S3): The PE-binding mutant, 34L88E-rhTIM1, more efficiently mediated ZIKV and WNV VLP infection than did WT-rhTIM1.

**Fig. 5.**
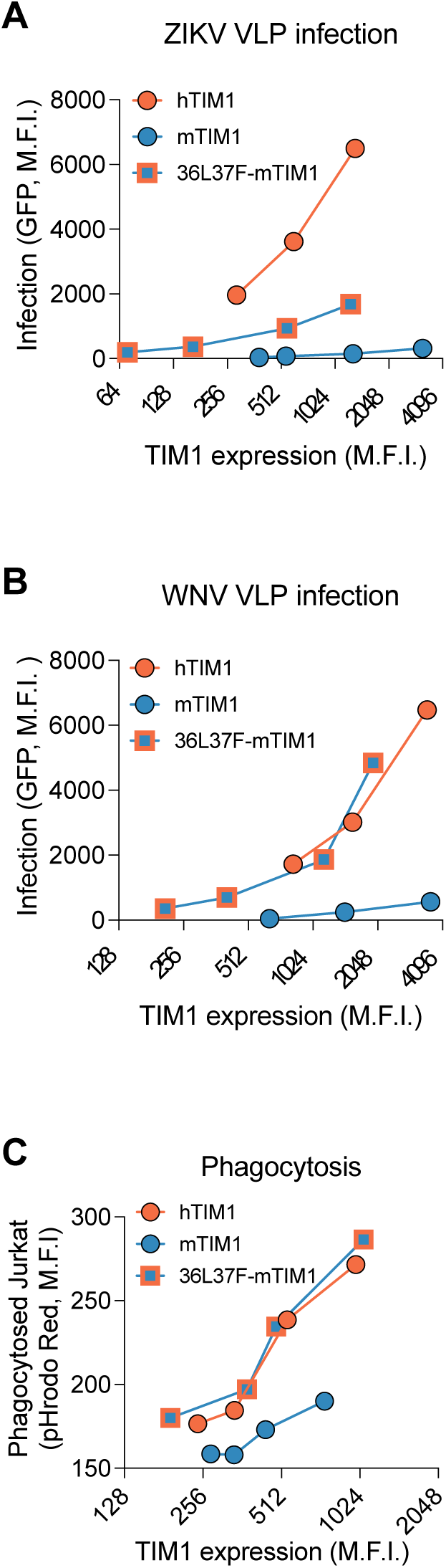
A PE-binding mutant of mTIM1 more efficiently mediates virus entry and efferocytosis. **(A) and (B)** VLP infection of 293T cells expressing WT-mTIM1 or PE-binding mTIM1 variant. 293T cells were transduced with serially diluted transducing vectors to achieve a wide range of expression levels of the indicated TIM1, MYC-tagged at their N-terminus. Next day, cells were replated on 6 and 48 well plates for staining and infection, respectively. Forty hours post transduction, the cells on 6 wells were assessed for TIM1 expression, using a MYC-tag antibody, and those on 48 wells were infected with ZIKV VLP (A) or WNV VLP (B). Infected cells were analyzed for GFP expression at 24 h post infection. **(C)** Phagocytosis of the apoptotic Jurkat cells mediated by WT- or PE-binding mTIM1 variant. Efferocytosis assays were performed in the same way as described in Fig. 1E except that 293T cells were transduced with varying amounts of vectors to obtain a wide range of TIM1 expression levels. Because statistical analysis is not possible in the presented data format, in which one variable (expression level) is not the same between the two groups (WT- and 36L37F-mTIM1), the results from two additional experiments each for A-C are shown in Fig. S2.

The efficiency of the 36L37F-mTIM1 to mediate efferocytosis was also measured. HEK293T cells similarly transduced to express WT-mTIM1, 36L37F-mTIM1, or hTIM1 at a wide range of levels and replated the next day. The following day, cells on 48 wells were assessed for efferocytosis and those on 6 wells for expression level. 36L37F-mTIM1 exhibited greater ability to mediate efferocytosis compared to WT-mTIM1 (Fig. 5C and S2C). In fact, 36L37F-mTIM1 was as efficient as hTIM1 in mediating efferocytosis. Together, these data make clear that PE-binding ability of TIM1 molecules is important for efficiency of their functions.

### The mucin domain contributes to binding apoptotic cells and virions

Although PE-binding mutants of mTIM1 and rhTIM1 exhibited greater efficiency in supporting VLP infection compared to their WT counterpart, we noticed a different pattern between ZIKV and WNV. The efficiency of 36L37F-mTIM1 and 34L88E-rhTIM1 is comparable to that of hTIM1 in supporting WNV VLP infection, but not for ZIKV VLP infection (Fig. 5A vs 5B, Fig. S2A vs S2B, and Fig. S3A vs S3B). In addition, although the head domain alone binds the PL ligand, the mucin domain was also reported to be important for mediating efferocytosis (10, 21). Thus, we compared the TIM1 head domain and the ectodomain, which contains the mucin stalk as well as the head domain, to assess the contribution of the mucin domain to VLP entry and efferocytosis. We first compared TIM1(head)-Fc_mono_ and TIM1(ecto)-Fc_mono_ for their ability to bind the flavivirus virions. Live ZIKV or WNV particles were incubated with Fc_mono_-fusion forms of TIM1(head) or TIM1(ecto) and precipitated with Protein A-Sepharose. Captured viral particles were either quantified by RT-qPCR (Fig. 6A) or visualized by Western Blot (WB) analyses using an antibody specific for ZIKV or WNV E protein (Fig. 6B). Both RT-qPCR and WB data demonstrate that substantially more ZIKV and WNV particles were captured by the ectodomain than by the head domain for all three TIM1 orthologs, although the difference was most prominent with hTIM1. To determine the contribution of the mucin domain in efferocytosis, we also compared the ability of the head and ecto domains to bind apoptotic Jurkat cells. As Fig. 6C and 6D show, TIM1(ecto)-Fc_mono_ exhibited much higher binding to the apoptotic Jurkat cells than did TIM1(head)-Fc_mono_ for all three TIM1 orthologs.

**Fig. 6.**
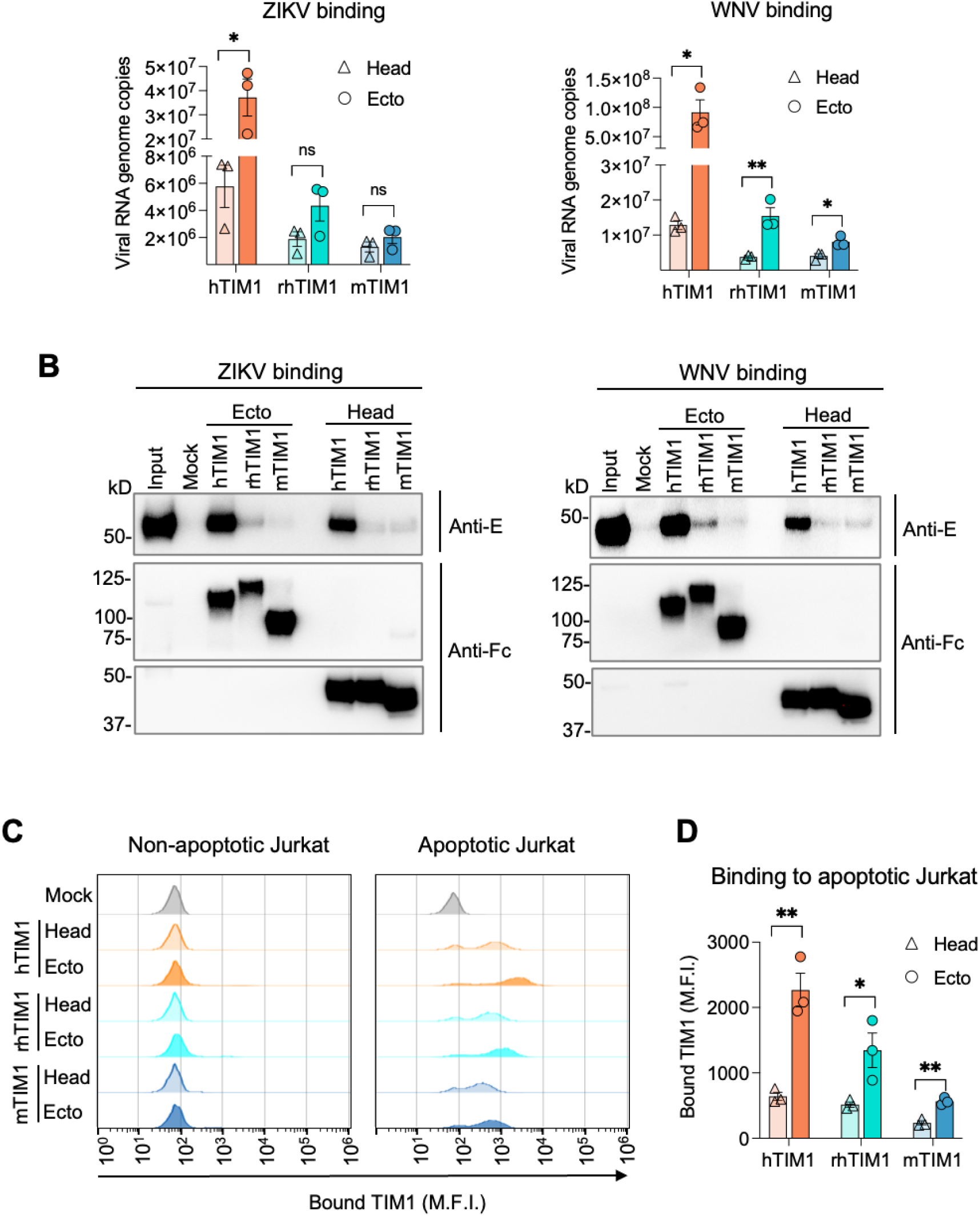
TIM1 mucin domain contributes to binding virions and apoptotic cells. **(A) and (B)** Pulldown assays of live ZIKV or WNV with the TIM1 head or ecto domain. Live virus particles (2x10^8^ genome copies) were incubated with 10 nM TIM1(head)-Fc_(mono)_ or TIM1(ecto)-Fc_(mono)_ protein in the presence of 2 mM Ca^2+^. Bound viruses were precipitated by the Protein A-Sepharose beads and analyzed either by RT-qPCR (A) or by WB (B). The sequences of qPCR primers and probe are provided in Table 1. For WBs, an aliquot of captured viruses was analyzed by non-reducing SDS-PAGE, and the E protein was detected using the pan-flavivirus antibody, 4G2 (top panels). The other aliquot was analyzed by reducing SDS-PAGE, and TIM1-Fc proteins were detected using an anti-human IgG antibody (bottom two panels). Each symbol in the data shown in (A) indicates the result form one independent experiment. WB images are the representatives from two independent experiments. **(C)** Binding of the apoptotic Jurkat cells by the TIM1 head or ecto domain. Actinomycin D (apoptotic) or DMSO (non-apoptotic) treated Jurkat cells were incubated with 2.5 nM TIM1(head)-Fc_(mono)_ or TIM1(ecto)-Fc_(mono)_ protein for 30 min at room temperature. Shown histograms are the representatives of three independent experiments. **(D)** Quantification of bound TIM1 shown in (C) and two additional experiments. (A and D) Data are presented as Mean ± SEM. Statistical significance was analyzed by unpaired t test. *p < 0.05, **p < 0.01; ns, not significant.

To make sure that the mucin domain did not alter PL ligand binding efficiency of the head domain, we compared PE and PS binding by the ectodomains to that of the head domain. The PE and PS binding profiles of TIM1(ecto)-Fc_mono_ and TIM1(head)-Fc_mono_, shown in Fig. S4 and Fig. 2, respectively, are nearly identical for all three TIM1 orthologs. These results together demonstrate that although the mucin domain does not aid the head domain in binding the PL ligands, it nonetheless enhances the efficiency of binding virions and apoptotic bodies in all three TIM1 orthologs.

### hTIM1 mucin domain cooperates with the PE-binding ability of the head domain in mediating virus entry and efferocytosis

After assessing the role of the mucin domain, using VLP and apoptotic body binding assays, we next investigated its contribution in VLP infection and efferocytosis. We compared the mucin domains of hTIM1 and mTIM1 by replacing the mucin domain of WT-mTIM1 and 36L37F-mTIM1 with that of hTIM1 (Fig. 7A) and assessing the resulting constructs for their ability to mediate flavivirus VLP entry and efferocytosis. Because the expression levels of WT and mutant TIM1 are widely different, we again conducted virus entry and efferocytosis assays at multiple different expression levels of WT and mutant TIM1. We observed that mTIM1-hMucin, which has hTIM1 mucin domain and WT mTIM1 head domain, did not enhance VLP entry compared to WT-mTIM1 (Fig. 7B, 7D, S5A, and S5C). Surprisingly, however, 36L37F-mTIM1-hMucin, the PE-binding mutant containing hTIM1 mucin domain clearly enhanced VLP entry (Fig. 7C, 7E, S5B, and S5D). This phenomenon was observed with both ZIKV and WNV VLPs, but a larger difference was noticed with ZIKV VLP.

**Fig. 7.**
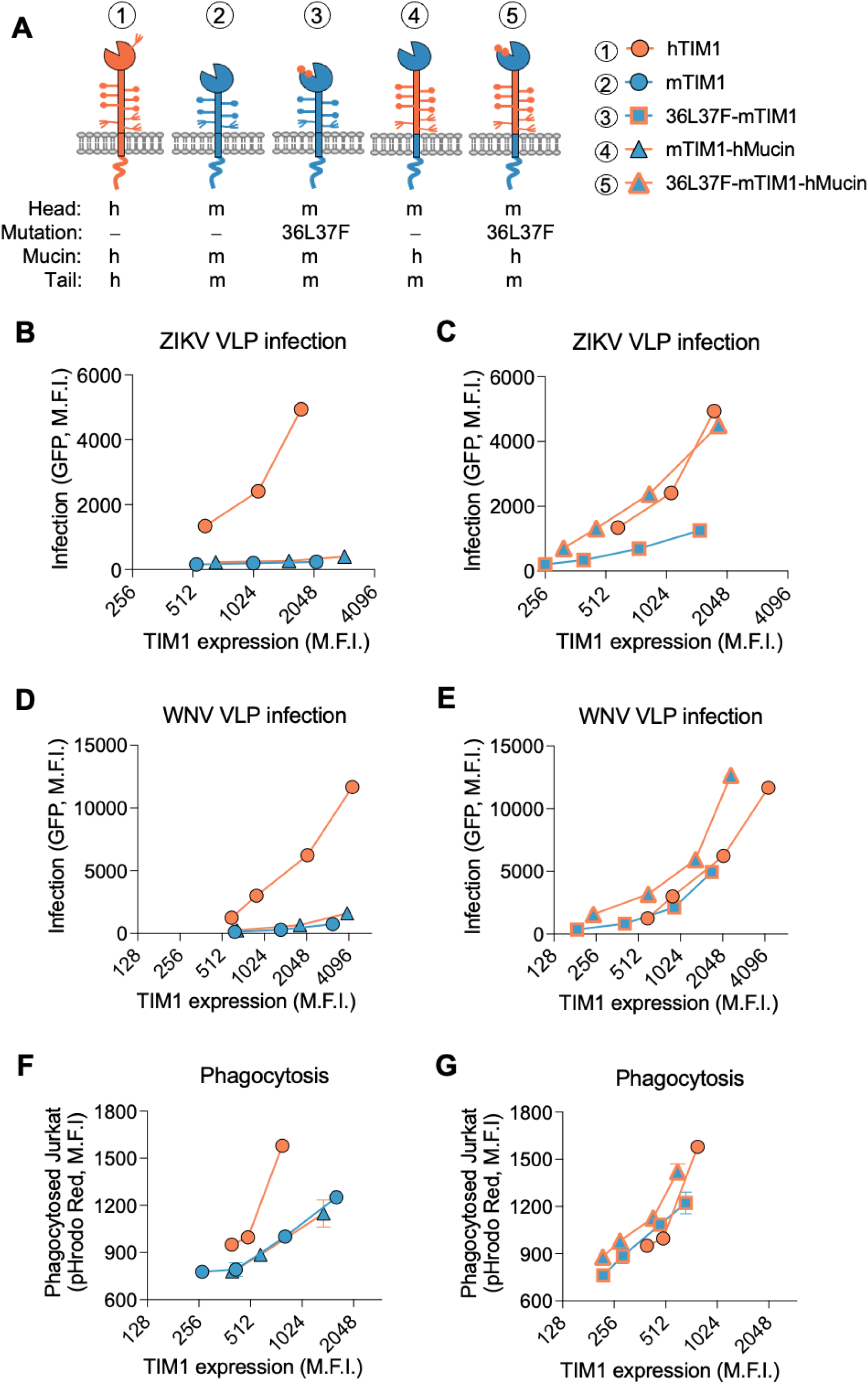
hTIM1 mucin domain cooperates with the PE-binding ability of the head domain in mediating virus entry and efferocytosis. **(A)** Diagrams of hTIM1, mTIM1, and mTIM1 variants used in (B-G), which include the PE-binding mutation (36L37F) and/or hTIM1 mucin domain (hMucin). The letters “m” and “h” indicate mouse and human, respectively. **(B-E)** VLP infection mediated by mTIM1 variants containing hTIM1 mucin domain. 293T cells expressing the indicated TIM1 molecules were infected with ZIKV VLP (B and C) or WNV VLP (D and E) in the same way as described in Fig. 5A and 5B. To help understanding, data on the five molecules are split into two groups: The data for hMucin in the presence of the WT-mTIM1 head domain (mTIM1 vs mTIM1-hMucin in B and D) and those in the presence of the PE-binding mTIM1 head domain (36L37F-mTIM1 vs 36L37F-mTIM1-hMucin in C and E). The data on hTIM1was used for both groups. **(F) and (G)** Phagocytosis of the apoptotic Jurkat cells by mTIM1 variants containing hTIM1 mucin domain. Efferocytosis assays were performed in the same way as described in Fig. 5C. Like in B-D, data are split in two groups to increase clarity: The effect of hTIM1 mucin domain in the presence of WT-mTIM1 head domain (F) and those in the presence of the PE-binding mTIM1 head domain (G). The data on hTIM1 was used in both groups. Because statistical analysis is not possible in the data format used in B-G, two additional experiments each for VLP infection and efferocytosis, performed in the same way, are shown in Fig. S5.

Regarding efferocytosis, although PE-binding mutation (36L37F) alone was sufficient to enhance efferocytosis efficiency of mTIM1 to that of hTIM1 (Fig. 5C and S2C), we nonetheless evaluated the effect of hTIM1 mucin domain in the background of WT- and 36L37F-mTIM1. Similar results as in VLP entry assays were obtained from efferocytosis assays: hTIM1 mucin domain did not have any effect when the head domain binds only PS (mTIM1-hMucin in Fig. 7F and S5E) but enhanced efferocytosis efficiency when the head domain has PE-binding ability, exceeding the efficiency of hTIM1 (36L37F-mTIM1-hMucin in Fig. 7G and S5F). These data demonstrate that superior performance of hTIM1 compared to other animal TIM1 orthologs is the result not only of the PE-binding ability of the head domain but also of the mucin domain that cooperates with its PE-binding head domain.

## DISCUSSION

We and others previously reported that hTIM1 can bind PE (10, 25) in addition to PS and that this PE-binding ability augments hTIM1’s role in mediating efferocytosis and virus uptake (25). In the current study, we show that this property is not shared by TIM1 orthologs of other species frequently used to model human diseases: Specifically, mTIM1 and rhTIM1 bind PS but not PE. We also show that PE-binding ability can be gained by mTIM1 and rhTIM1 by replacing two of their residues in the head domain with the hTIM1 equivalents. Further, consistent with previous reports (21), we found that the mucin domain also contributes to TIM1’s activities to mediate efferocytosis and support virus entry. Collectively, these observations demonstrate that hTIM1 is more active than mTIM1 and rhTIM1 and imply that TIM1 studies in animal models may not fully describe the extent of hTIM1 functions. One question that arises is why only hTIM1 has the unique ability to bind PE. Although we do not currently know the answer, what we know is that the proteins that bind only PS still benefit from the presence of PE in the membrane, owing to the synergy between PE and PS (56–58). For example, like mouse and rhesus TIM1, human TIM4 and GAS6 bind PS but not PE. However, when PE and PS are present together, which is the case for most biological membranes, TIM4 and GAS6 binding to PS is drastically increased without gaining PE binding ability. Therefore, although most PS-binding proteins do not directly bind PE, they nonetheless benefit from the presence of PE in the membrane.

A number of groups highlighted the role of the mucin domain in TIM1’s functions. Briefly, upon kidney injury in healthy mice, TIM1 expression was induced in the tubular epithelial cells to clear neighboring apoptotic cells, but mice in which TIM1 mucin domain was genetically deleted could not clear the apoptotic cells (10, 21). Further, mice in which TIM1 mucin domain is deleted exhibited defective regulatory B cell functions and developed spontaneous autoimmunity when they aged, indicating contribution of TIM1 mucin domain to immune regulation (15). Consistent with these reports, we also found the mucin domain contributes to the functions of all three TIM1 orthologs (Fig. 7 and S5). Notably, we observed that hTIM1 mucin domain displayed enhanced function only in the presence of a head domain that is able to bind PE in addition to PS. This is unexpected because the mucin domain does not directly bind PE or promote PE association (Fig. 2 and S4). Nevertheless, the ability to phagocytose apoptotic cells and to support VLP infection was enhanced when hTIM1 mucin domain was introduced to mTIM1 with the PE-binding head domain (36L37F-mTIM1-hMucin, Fig 7C, E, and G) but not when it was introduced to mTIM1 with the WT head domain (mTIM1-hMucin, Fig. 7B, D, and F). It remains unclear how PE-binding head domain cooperates with the mucin domain, but there are a few potential mechanisms. First, greater energy might be provided by PS and PE binding, owing to the higher frequency with which TIM1 molecules come in contact with PE as well as PS compared to PS alone. Tighter association with the target membrane provided by binding both PE and PS could facilitate local interactions between the mucin domain and its protein or glycan binding partner present on the virions or apoptotic cells. Such association then in turn could help stabilize the interaction between the head domain and the virion or apoptotic membrane. Alternatively, it is also possible the binding partners of the mucin domain are preferentially localized in the membrane microdomains that also contain or are enriched with PE, and thus the mucin domain can more easily gain access to those binding partners when the head domain binds PE.

In summary, our study here highlights the quantitative and qualitative differences between hTIM1 and its rhesus macaque and mouse orthologs in both the head and mucin domains. Thus, the studies of TIM1 that relies on mouse or macaque models to draw conclusions about human physiology may require additional caveats.

## MATERIALS AND METHODS

### Cell lines

Human embryonic kidney HEK293T cells were grown in high-glucose DMEM (Life Technologies, Cat# 10569-010), and Jurkat (human T lymphocyte) cells in RPMI 1640 medium (Life Technologies, Cat# 61870-036). All cells were cultured in medium supplemented with 10% FBS (Sigma-Aldrich, Cat# F2442) and 100 U/mL each Penicillin and Streptomycin (Life Technologies, Cat# 15140-122) at 37°C with 5% CO_2_. 293T cells transduced to stably express TIM1 (TIM1-293T) or mock transduced (Mock-293T) were maintained in the medium supplemented with 1 μg/mL puromycin (InvivoGen, Cat# ant-pr).

### Pseudovirus (PV) and Virus Like Particle (VLP) production

The expression plasmids encoding the entry glycoprotein of Zaire Ebola virus (EBOV, Mayinga strain), Eastern Equine Encephalitis virus (EEEV, FL91-4697 strain), Vesicular Stomatitis virus (VSV, Indiana strain), Lassa fever virus (LASV, Josiah stain), Zika virus (ZIKV, Brazil strain) and West Nile virus (WNV, NY99 stain) were previously described (25, 39, 56). The retroviral vector expressing enhanced green fluorescence protein (eGFP), pQCXIX-eGFP, and the WNV replicon expressing eGFP were also described in the previous studies (25, 39, 56).

To produce murine leukemia virus (MLV) based PVs for making TIM1-expressing stable cells, the pQCXIP-TIM1s were transfected into 293T together with a plasmid encoding the MLV gag-pol protein and a plasmid encoding the VSV G protein. The PV for mock transduction was produced using empty pQCXIP plasmid.

Similarly, PVs bearing various viral entry glycoproteins were produced in 293T cells as described previously (56) by transfection of pQCXIX-eGFP together with two plasmids separately encoding MLV gag-pol and a viral entry glycoprotein. The genes for viral entry glycoproteins are described above. PVs were harvested from the cell culture supernatants at 32-34 h post-transfection. To produce ZIKV and WNV VLPs, 293T cells were transfected with a plasmid encoding WNV replicon (59) and a plasmid encoding either WNV-C/ZIKV-prME or WNV-CprME (59) at a 2:1 ratio. The plasmid expressing WNV-C/ZIKV-prME was generated by replacing WNV-prME with that of ZIKV (Brazil strain). Cell culture supernatants containing VLPs were harvested at 48 h post-transfection. Both the supernatants containing PVs and VLPs were clarified by 0.45 μm filtration and aliquoted for storage at -80°C before use.

### Wild-type and mutant TIM1-Fc_(mono)_ protein production

The expression plasmids for TIM1(head)-Fc_(mono)_ and TIM1(ecto)-Fc_(mono)_ fusion proteins were constructed by cloning the coding sequences of the head domain (residues 21-126 for both hTIM1 and rhTIM1 and 22-129 for mTIM1) or the ecto domain (residues 21-290 for hTIM1, 21-355 for rhTIM1, and 22-237 for mTIM1) into pcDNA3.1 (+) vector containing the CD5 signal peptide and the genomic sequence of the human IgG1 Fc region. To prevent dimerization three mutations (L368R, F495H, and Y407E) as well as the mutations of three cysteines involved in interchain disulfide bonds (C310A, C316N and C319G) were introduced into the Fc domain (55). The mutant and chimeric TIM1 constructs were made by overlapping PCR.

To produce TIM1(head)-Fc_(mono)_ or TIM1(ecto)-Fc_(mono)_ proteins, 293T cells were transfected with an appropriate plasmid by calcium-phosphate and cultured in FreeStyle 293 medium (Thermo Fisher, Cat# 12338018) for 72 h. The culture supernatants were harvested and clarified by 0.45 μm filtration. These proteins were precipitated by Protein A Sepharose beads for the quantification by Coomassie blue staining following a non-reducing SDS-PAGE. Purified human IgG was used as a standard.

### TIM1 stable cell line construction

The plasmids expressing the full-length human TIM1 (GenBank: AAC39862.1), rhesus TIM1 (GenBank: OR896543), and mouse TIM1 (GenBank: NP_599009.2) were generated by cloning their corresponding cDNA fragments into the retroviral vector, pQCXIP (Clontech). All TIM1 orthologs used in this study are expressed with the signal peptide of mouse angiotensin-converting enzyme 2 (MSSSSWLLLSLVAVTTAQ) and the MYC-tag (EQKLISEEDL) at their N-termini.

To make 293T cells stably expressing the wild-type full-length TIM1 from human, mouse, and rhesus, 30-40% confluent 293T cells were seeded in 6-well plate and transduced with the PV expressing the indicated TIM1. The PVs used to make TIM1-expressing stable cells, were produced by transfecting 293T cells with pQCXIP-TIM1 plasmid with a plasmid encoding the MLV gag-pol protein and that encoding the VSV G protein. To produce Mock-293T, cells were transduced with the PV produced using the empty pQCXIP plasmid. Two days later, the transduced cells were selected with 1 μg/mL puromycin. The stable TIM1-293Ts and Mock-293T were maintained in 1 μg/mL puromycin in culture.

### Cell surface staining for TIM1

To determine cell surface expression level of different TIM1 molecules on either stably or transiently transduced 293T cells, the cells were detached with 5mM EDTA in PBS, washed, and stained with 3 μg/mL MYC-tag antibody (clone 9E10) in PBS containing 2% goat serum followed by 2 μg/mL goat anti-mouse IgG conjugated with Alexa 647 (Jackson ImmunoResearch, Cat# 115-606-146). 9E10 antibody was purified using Protein A-Sepharose beads from the culture supernatant of the hybridoma cell line (CRL-1729) purchased from American Type culture collection. Washed cells were read by Attune NxT flow cytometer equipped with an autosampler CytKick (Thermo Fisher), and the data were analyzed using FlowJo (FlowJo, LLC).

### PV and VLP entry assay

To assess viral entry efficiency mediated by various TIM1 molecules and their variants, 293T cells expressing these proteins were seeded 24 hours prior to infection at 3x10^4^ cells/well in the 48-well plates coated with 0.1 mg/mL poly-D-lysine (Sigma-Aldrich, Cat# P6403). The same cells were also plated on the 6-well plates to assess TIM1 expression level. Next day, the cells on the 48-well plates were incubated either with PVs or with flavivirus VLPs. After 1 h infection at 37°C, cells were replenished with the fresh medium after removing the PVs or VLPs. At the time of infection, the cells on the 6-well plates were detached using 5 mM EDTA in PBS and stained with the anti-MYC tag antibody (9E10) for the measurement of surface TIM1 expression level. Infected cells were harvested, and GFP was read by flow cytometry at 24 h post-infection.

### Live virus infection assay

To further confirm the viral entry efficiency mediated by TIM1s, replication-competent ZIKV (PB81 strain) grown in Vero cells were used to infect 293T cells that stably express TIM1s. Cells were infected with varying amounts of virus at 37°C. Virus was removed 1 h later and cells were replenished with fresh culture media. At 24 h post-infection, cells were harvested by trypsinization, fixed with 2% paraformaldehyde in PBS, and permeabilized with 0.05% Saponin in PBS for intracellular staining of the Envelope (E) protein with 1 μg/mL pan-flavivirus antibody, 4G2, followed by 2 μg/mL goat anti-mouse IgG (H+L) conjugated with Alexa 647 (Jackson ImmunoResearch, Cat# 115-606-146). 4G2 antibody was purified using Protein A-Sepharose from the culture supernatant of the hybridoma cell line (HB-112) purchased from American Type Culture Collection. Washed cells were read by Attune NxT flow cytometer equipped with an autosampler CytKick (Thermo Fisher), and the data were analyzed using FlowJo (FlowJo, LLC).

### Efferocytosis assay

To investigate the TIM1-mediated phagocytosis of apoptotic cells, 293T cells expressing TIM1 were seeded in a 48-well plate at 1 x 10^4^ cells per well one day before the assay. In parallel, Jurkat cells (2.5 x 10^5^/mL) were treated with 1 μM Actinomycin D (Thermo Fisher, Cat# A7592) in 10 mL complete RPMI media in T25 Flask for 15 hours in 5% CO2 incubator to induce apoptosis. Jurkat cells treated with DMSO were included as a negative control. After treatment, the Jurkat cells were washed once with the wash buffer provided by pHrodo Red Cell Labeling Kit for Incucyte (Sartorius, Cat# 4649) and resuspended at 1 x 10^6^ cells/mL labeling buffer containing 0.1 μg/mL pHrodo Red dye and incubated for 1 h in a CO_2_ incubator. Labeled Jurkat cells were washed once with complete RPMI media and resuspended in complete DMEM media at 1 x 10^6^ cells/mL. Phagocytosis assay was performed by co-incubating labeled Jurkat cells with TIM1-expressing 293T cells at a 10:1 ratio (Jurkat : TIM1-293T) for 1 h at 37°C, while cells incubated on ice were included as a negative control. Unbound Jurkat cells were removed by PBS wash, and 293T cells were detached from the plate by trypsinization. Phagocytosis was assessed by measuring the pHrodo Red florescence by flow cytometry in the 293T cell gate.

### Phospholipids ELISA

The following phospholipids (Avanti Polar Lipids) were used in assays: 1,2-dioleoyl-sn-glycero-3-phosphocholine (PC, Cat# 850375), 1,2-dioleoyl-sn-glycero-3-phosphoethanolamine (PE, Cat# 850725), 1,2-dioleoyl-sn-glycero-3-phospho-L-serine (PS, Cat# 840035), 1,2-dioleoyl-sn-glycero-3-phospho-[1’-myo-inositol] (PI, Cat# 850149) and Sphingomyelin (SPH, Cat# 860062). As described in our previous study (25, 56), to assess phospholipid-binding profiles of TIM1-Fc_(mono)_ proteins, polystyrene ELISA plates (Falcon, Cat# 351172) were coated with the indicated amounts of phospholipids in methanol and dried out completely at room temperature overnight. The plates were washed with Tris-buffered saline (TBS: 25 mM Tris base, 137 mM NaCl, 2.7 mM KCl, pH 7.4) containing 2 mM CaCl_2_ and 0.05% (vol/vol) Tween 20 (TBST-Ca^2+^), blocked with 1% bovine serum albumin (BSA) in TBS for 1 h at room temperature, and washed three times with TBST-Ca^2+^. Then 1 nM TIM1-Fc_(mono)_ proteins in TBS containing 2 mM CaCl2 (TBS-Ca^2+^) were added, and the plates were gently rocked at room temperature for 1 h. The plates were washed three times with TBST-Ca^2+^ before incubating with a goat anti-human IgG conjugated with horseradish peroxidase (Jackson ImmunoResearch, Cat# 109-035-098). Bound TIM1-Fc_(mono)_ proteins were visualized using UltraTMB substrate (Thermo Fisher, Cat# 34028) after the plates were extensively washed with TBST-Ca^2+^ and TBS-Ca^2+^. Reaction was terminated with 2 M phosphoric acid, and the plates were read at 450 nm in a SpectraMax Paradigm microplate reader (Molecular Devices). Wells treated the same way but only coated with methanol were used as background controls.

### Live virus pull-down assay

To assess the binding affinity of TIM1-Fc_(mono)_ proteins to the live virus particles, 2x10^8^ genome copies of ZIKV or WNV were incubated with 10 nM TIM1-Fc _(mono)_ proteins in 500 μL TBS-Ca^2+^ at 37°C for 1 h followed by the incubation with 20 μL of 50% (vol/vol) protein A-Sepharose beads by rocking at room temperature for another 1 h. Beads were spun down at 1000 x g for 3min and washed three times with TBST-Ca^2+^ to remove uncaptured viruses. Captured viruses were detected either by RT-qPCR or by Western-Blot (WB). For RT-qPCR, viral RNA was extracted from the precipitated beads using TRIzol (Invitrogen, Cat# 10296028) and GlycoBlue coprecipitant (Invitrogen, Cat# AM9516) and reverse transcribed using a high-capacity cDNA reverse transcription kit (Applied Biosystems, Cat# 4374966). qPCR was performed using Luna Universal Probe qPCR Master Mix (New England Biolabs, Cat# M3004) with specific primers and probes targeting NS3 gene of ZIKV or WNV (Table 1), synthesized by Integrated DNA Technologies, using the PCR protocol: 95°C for 3min x 1 cycle, 95°C for 5 sec and 60°C for 30 sec x 40 cycles. Known quantity of a plasmid containing the targeted NS3 gene fragment of ZIKV or WNV was used to generate standard curves. For WB, viruses captured by the beads were analyzed by nonreducing and reducing SDS-PAGE, transferred to polyvinylidene difluoride (PVDF) membranes, and blotted with the pan-flavivirus antibody, 4G2, to detect the E protein (non-reducing gels). An anti-human Fc antibody (Jackson ImmunoResearch, Cat# 109-035-098) was used to detect the TIM1-Fc_(mono)_ proteins on reducing gels. Bands were visualized using the SuperSignal West Atto ultimate-sensitivity substrate (Thermo Scientific, Cat# A38555) and images were captured by ChemiDoc (Bio-Rad).

**Table 1.**
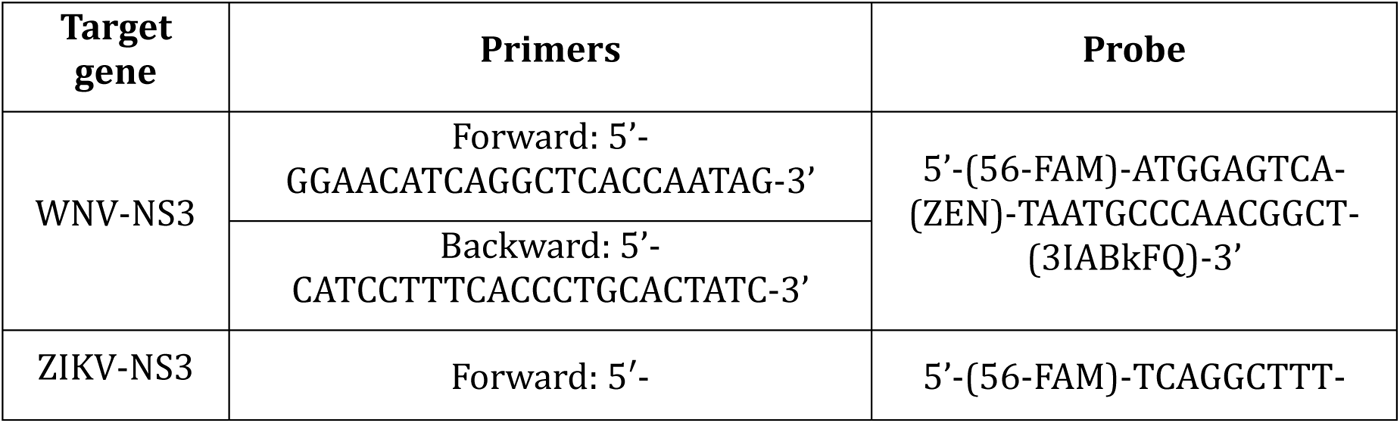
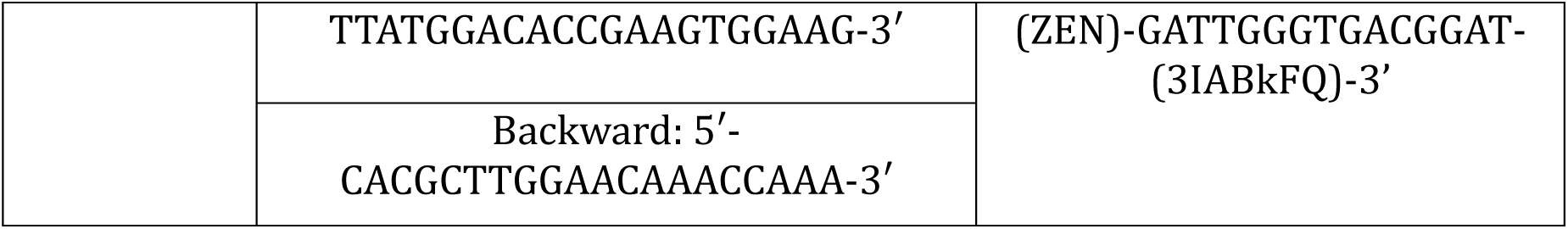
The sequences of primers and probe used for WNV and ZIKV qPCR.

### Binding assay of apoptotic Jurkat cells with TIM1-Fc proteins

To assess the binding affinity of TIM1-Fc_(mono)_ to apoptotic cells, Jurkat cells were induced to apoptose with 1 μM Actinomycin D as aforementioned. 5 x 10^4^ cells per well in a 96-well V-bottom plate were washed once with the binding buffer (10 mM HEPES, 140 mM NaCl, and 2.5 mM CaCl2) followed by incubation with 2.5 nM TIM1-Fc_(mono)_ proteins at room temperature for 30 min. After removing the unbound TIM1-Fc_(mono)_ proteins by washing once with the binding buffer, cells were incubated with 2 μg/mL goat anti-human IgG (H+L) Alexa647 (Jackson ImmunoResearch, Cat# 109-605-003) for 30 min on ice. Cells were read by flow cytometry (Attune, NxT, Thermo Fisher) after washing three times with the binding buffer, and the data analyzed by FloJo (FlowJo, LLC).

### Statistical analysis

All data was analyzed with GraphPad Prism version 9.0 (GraphPad Software Inc.) and expressed as Mean ± standard error of the mean (SEM). The difference within the group was tested by unpaired or paired t test, while between groups was tested using either one-way or two-way analysis of variance (ANOVA). Specific statistical analysis methods are described in the figure legends where results are presented. Values are considered statistically significant for *p* < 0.05.

## Data, Materials, and Software Availability

All study data are included in the article and/or SI Appendix.

## ACKNOWLEDGMENTS

We thank Drs. Joseph V. Bonventre (Brigham and Women’s Hospital, Harvard Medical School, Boston, MA), Takaharu Ichimura (Brigham and Women’s Hospital, Harvard Medical School, Boston, MA), and Vijay K. Kuchroo (Broad Institute, Brigham and Women’s Hospital, Mass General Hospital, and Harvard Medical School, Boston, MA) for their critical reading of the manuscript and insightful suggestions. This work was supported by the NIH grant R01 AI110692 to H.C.

## AUTHOR CONTRIBUTIONS

L.Z. and H.C. designed research; L.Z., C.E. K., A.S.R., and S.P. performed research; L.Z., C.E.K., and H.C. analyzed data; L.Z. and H.C. wrote the paper.

## SUPPLEMENTAL FIGURES

**Fig. S1 (Related to Fig. 1).**
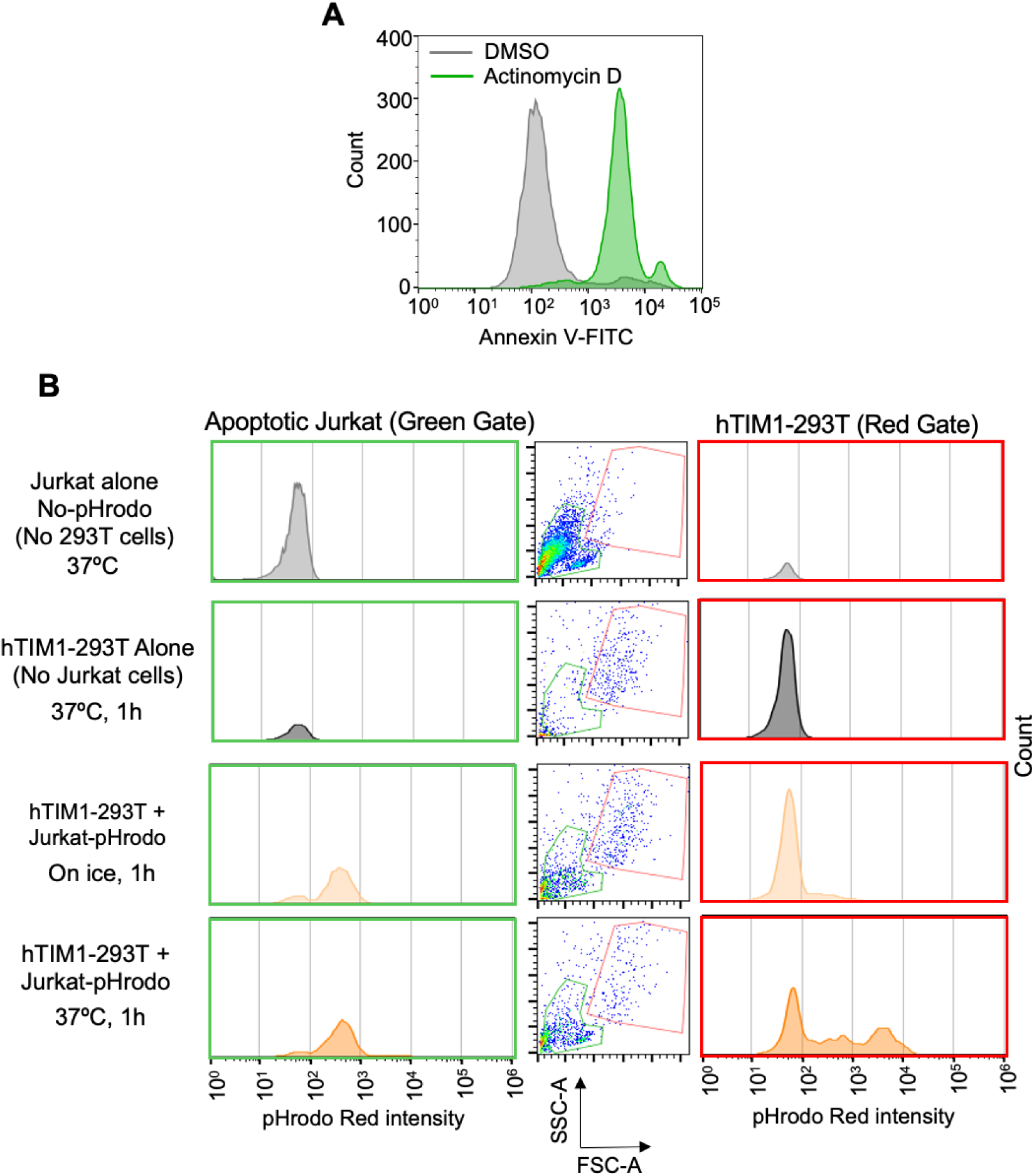
Phagocytosis of apoptotic Jurkat cells by hTIM1-293T. **(A)** Apoptosis induction in Jurkat cells. Cells were treated with 1 μM Actinomycin or DMSO for 15 hours and stained with Annexin V to detect apoptotic cells. **(B)** Gating strategy for phagocytosed Jurkat cells. Actinomycin D or DMSO treated Jurkat cells were loaded with 0.1 μM pHrodo Red in a CO_2_ incubator for 1 h, added to Mock-or TIM1-293T for 1 h incubation. Following washing and detaching 293T cells via trypsinization, 293T cells and uninternalized Jurkat cells were gated based on samples containing 293T or Jurkat cells alone. Of the three columns, the panels in the left column show fluorescence from free (unbound or unwashed) Jurkat cells, those in the middle column show the gates for Jurkat (green) and 293T (red) cells, and those in the right column show the fluorescence from 293T cells harboring phagocytosed Jurkat cells. The histogram in the 3rd row of the right-side column (co-incubation on ice) demonstrates that only minimum florescence is emitted from the uninternalized Jurkat cells that nonetheless are attached to 293T cells. The histogram in the 4th row of the right-side column (co-incubation at 37°C) demonstrates that intense red signal is emitted from Jurkat cells only if phagocytosis is allowed at 37°C.

**Fig. S2 (Related to Fig. 5).**
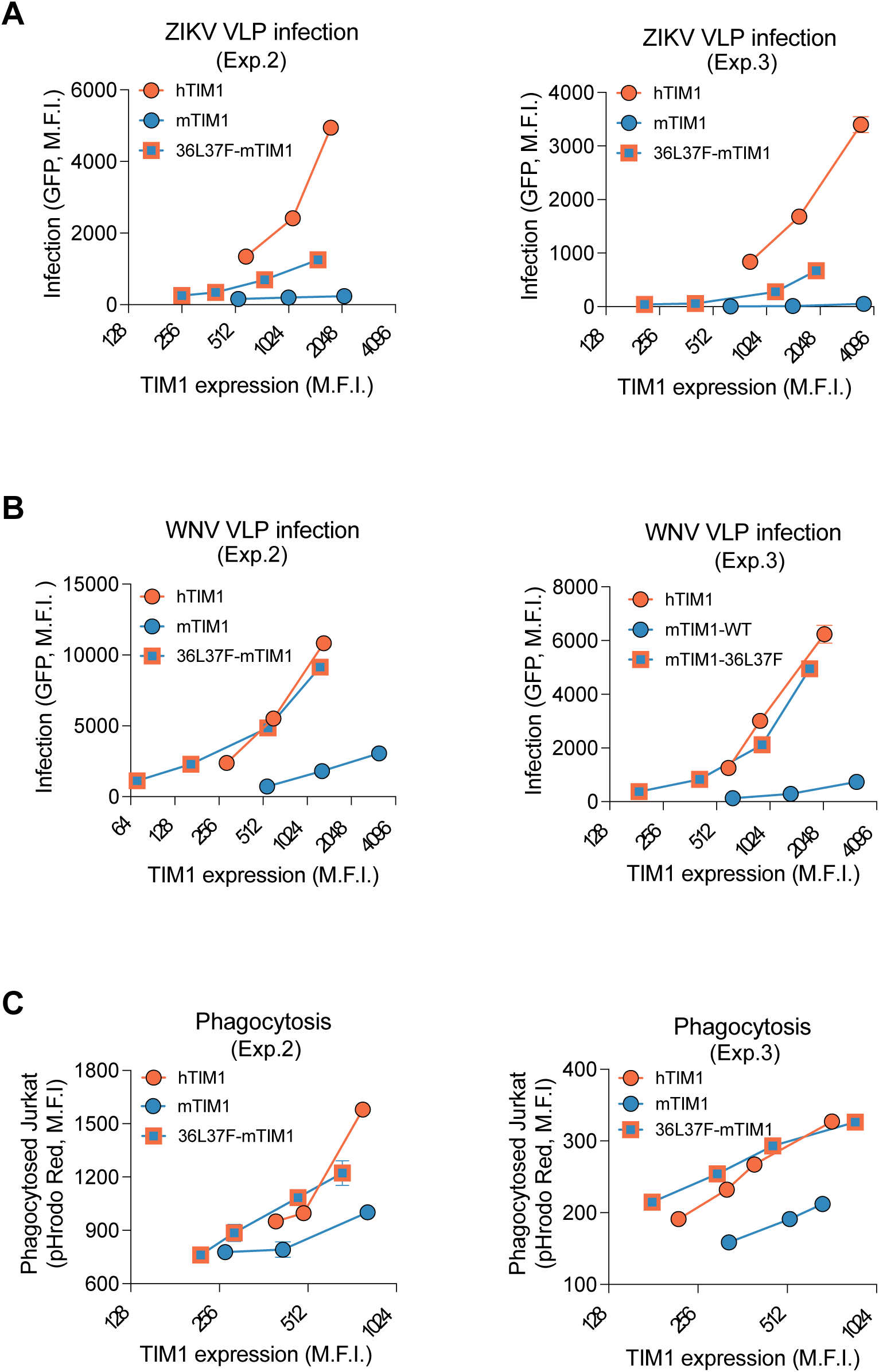
A PE-binding mutant of mTIM1 more efficiently mediates virus entry and efferocytosis. **(A) and (B)** Two additional experiments for infection by ZIKV VLP (A) or WNV VLP (B) mediated by PE-binding mTIM1 mutant, 36L37F-mTIM. **(C)** Two additional efferocytosis assays mediated by PE-binding mTIM1 mutant. VLP infection and efferocytosis assays were performed in the same way as described in Fig. 5.

**Fig. S3 (Related to Fig. 5).**
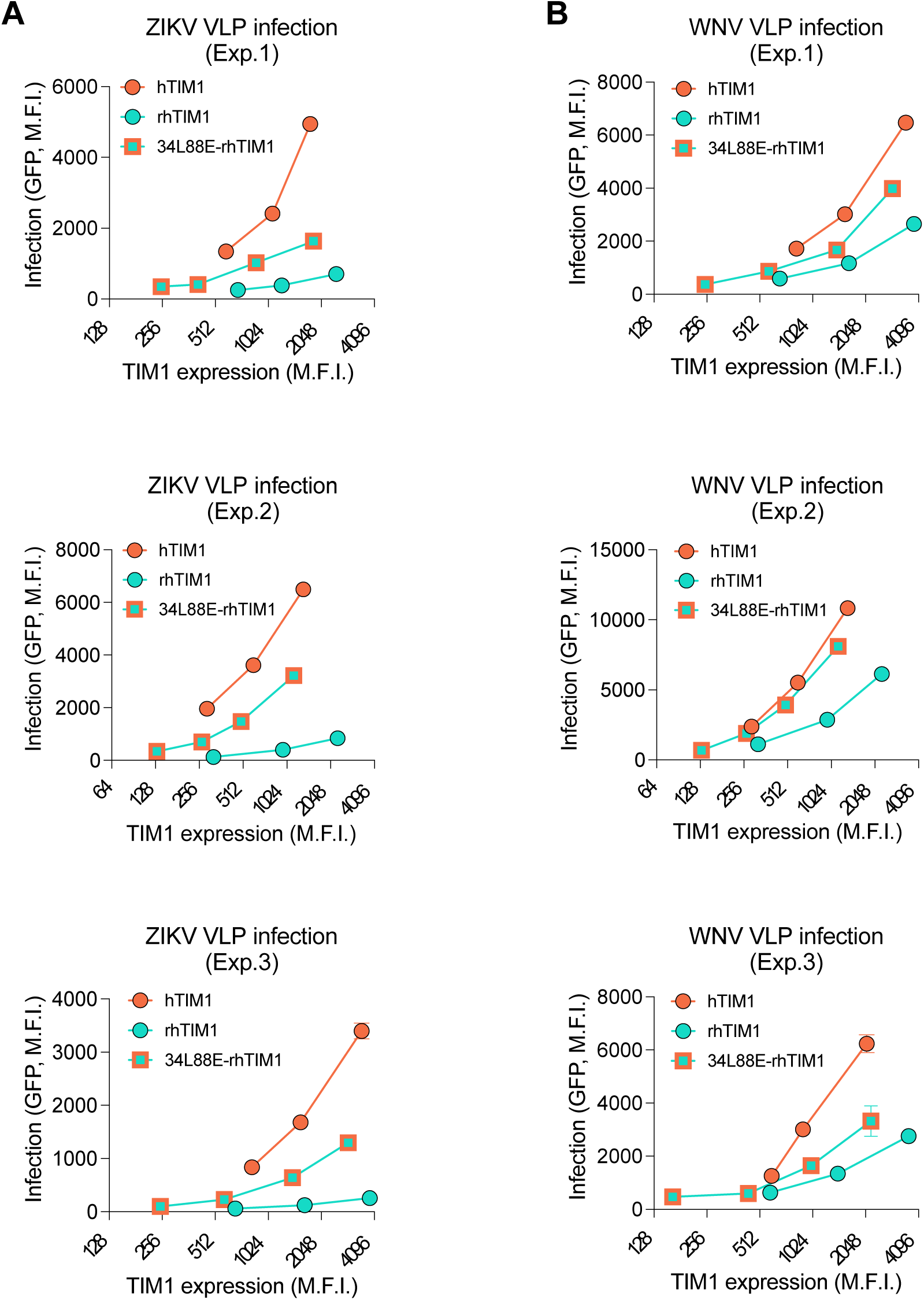
A PE-binding mutant of rhTIM1 more efficiently mediates virus entry. Three independent infection experiments using ZIKV VLP **(A)** or WNV VLP **(B)** were performed in the same way as described in Fig. 5A and B except that PE-binding rhTIM1 mutant (34L88E-rhTIM1) was compared to rhTIM1 and hTIM1.

**Fig. S4 (Related to Fig. 6).**
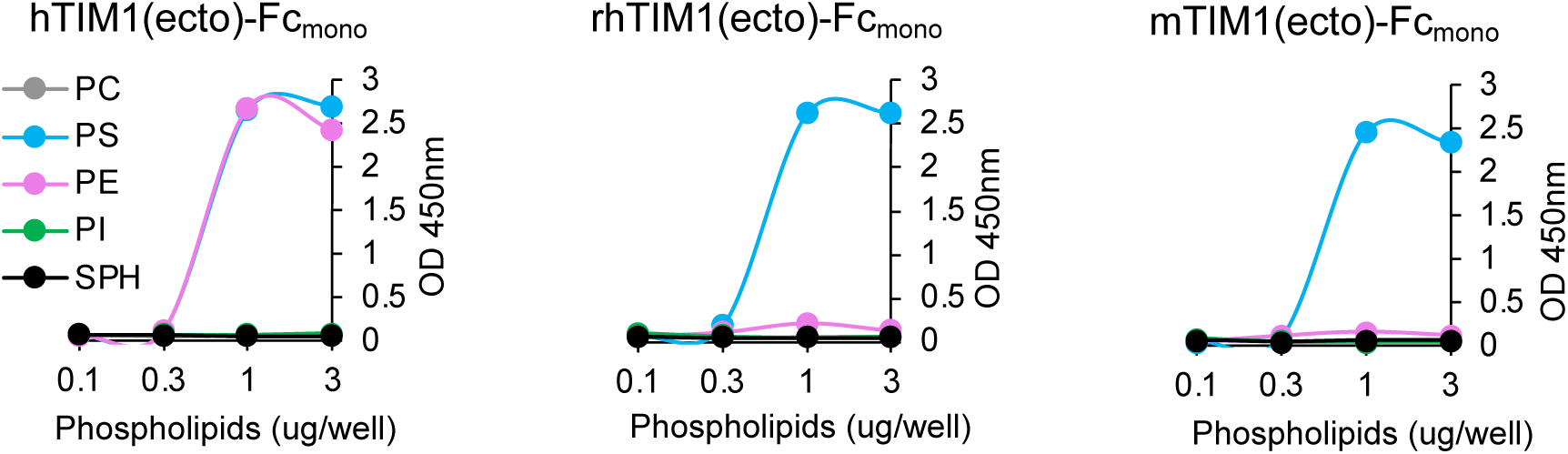
Phospholipid binding profiles of TIM1 ectodomain derived from three TIM1 orthologs. Increasing amounts (0.01 to 3 μg per well) of the indicated phospholipids dissolved in methanol was completely air dried on ELISA plates. Plates were washed with 0.05% TBST, blocked with 1% BSA, and incubated for 1 h at room temperature with 100 μL of 1 nM TIM1(ecto)-hFc_(mono)_ protein in TBS containing 2 mM Ca^2+^. The data shown here are the representatives of three independent experiments with similar results.

**Fig. S5 (Related to Fig. 7).**
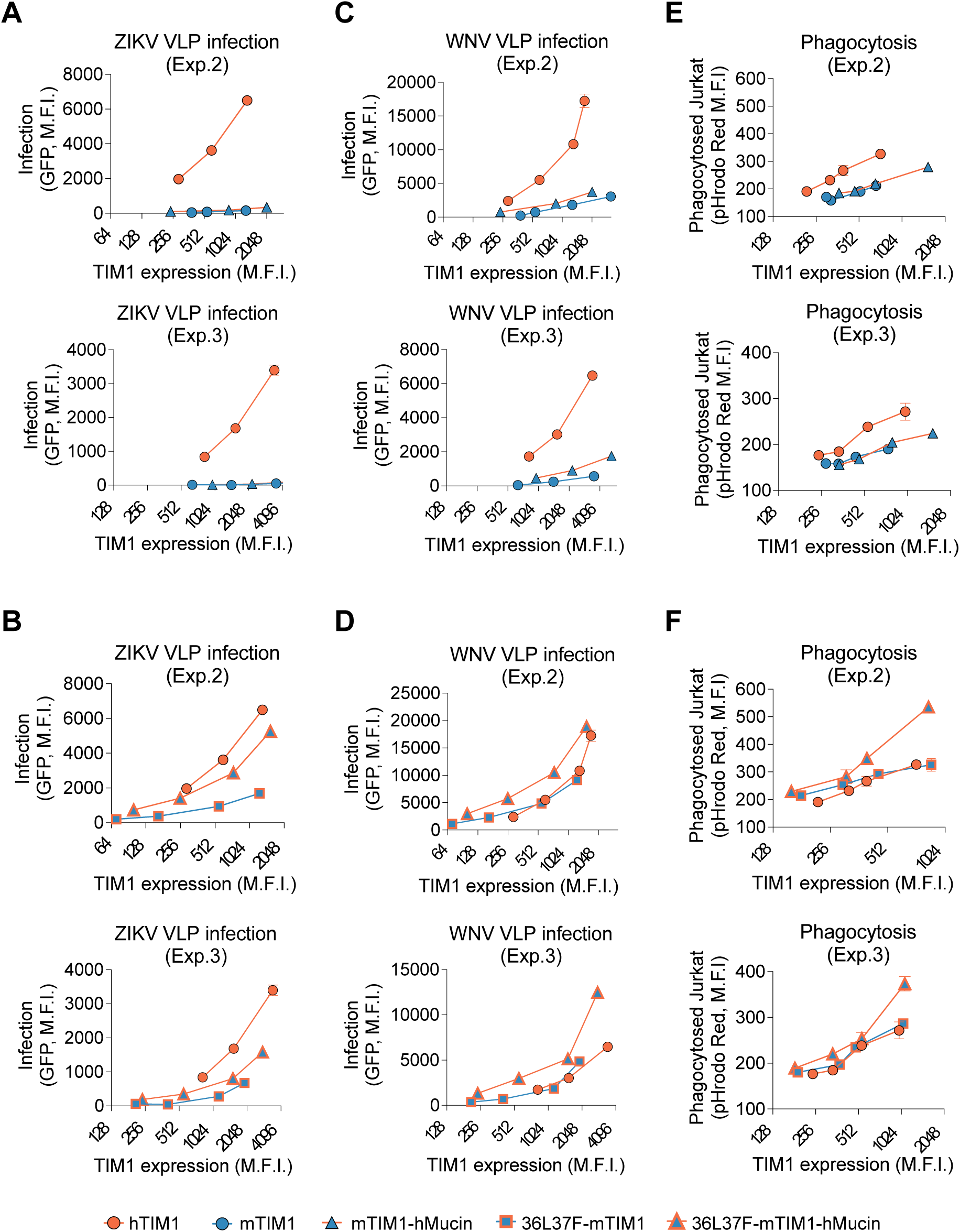
hTIM1 mucin domain cooperates with the PE-binding ability of the head domain in mediating virus entry and efferocytosis. **(A-D)** Two additional infection experiments for ZIKV VLP (A and B) and WNV VLP (C and D) mediated by mTIM1 with or without containing the hTIM1 mucin domain (A and C) or mediated by 36L37F-mTIM1 with or without containing the hTIM1 mucin domain (B and D). Experiments were performed as described in Fig. 7B-E. **(E) and (F)** Two additional experiments of efferocytosis mediated by mTIM1 with or without containing the hTIM1 mucin domain (E) or mediated by 36L37F-mTIM1 with or without containing the hTIM1 mucin domain (F). Experiments were performed as described in Fig. 7F and G.

